# TUGDA: Task uncertainty guided domain adaptation for robust generalization of cancer drug response prediction from *in vitro* to *in vivo* settings

**DOI:** 10.1101/2020.12.17.415737

**Authors:** Rafael Peres da Silva, Chayaporn Suphavilai, Niranjan Nagarajan

## Abstract

Over the last decade, large-scale cancer omics studies have highlighted the diversity of patient molecular profiles and the importance of leveraging this information to deliver the right drug to the right patient at the right time. Key challenges in learning predictive models for this include the high-dimensionality of omics data, limitations in the number of data points available, and heterogeneity in biological and clinical factors affecting patient response. The use of multi-task learning (MTL) techniques has been widely explored to address dataset limitations for *in vitro* drug response models, while domain adaptation (DA) has been employed to extend them to predict in vivo response. In both of these transfer learning settings, noisy data for some tasks (or domains) can substantially reduce the performance for others compared to single-task (domain) learners, i.e. lead to negative transfer (NT). We describe a novel multi-task unsupervised domain adaptation method (TUGDA) that addresses these limitations in a unified framework by quantifying uncertainty in predictors and weighting their influence on shared domain/task feature representations. TUGDA’s ability to rely more on predictors with low-uncertainty allowed it to notably reduce cases of negative transfer for *in vitro* models (63% for drugs with limited data and 94% overall) compared to state-of-the-art methods. For domain adaptation to *in vivo* settings, TUGDA improved performance for 6 out of 12 drugs in patient-derived xenografts, and 7 out of 22 drugs in TCGA patient datasets, despite being trained in an unsupervised fashion. TUGDA’s ability to avoid negative transfer thus provides a key capability as we try to integrate diverse drug-response datasets to build consistent predictive models with *in vivo* utility.

**Availability:** https://github.com/CSB5/TUGDA

## 1 Introduction

Advances in DNA sequencing technologies have galvanized a paradigm shift in medicine from a one-size-fits-all approach to precision medicine, that is tailored to stratified populations based on molecular information [7]. In oncology, an appreciation of the molecular diversity of cancers and limitations of standard-of-care treatments have further driven this interest towards patient-specific options based on re-purposing drugs and identifying targeted drug combinations [6]. The availability of a large number of cancer cell lines has provided ready models for collecting drug response data [16]. In combination with detailed omics profiles, these datasets present a unique opportunity to advance precision oncology based on state-of-the-art machine learning techniques [18].

The complexity inherent in biological systems and omics data poses two main challenges in learning models that could have clinical utility. Firstly, the high-dimensionality of omics data relative to the number of data points available can impact the generalizability of the models that are learnt [4]. Joint models that predict response for many drugs in a multi-task learning (MTL) setting have been widely used to alleviate this limitation [38,8,35,32]. Secondly, while cell line datasets are typically used to learn predictive models, they are not expected to capture key aspects relevant to *in vivo* response including tumor heterogeneity and microenvironment, immune response and overall patient health [34]. To partially address this, some recent methods have sought to use domain adaptation (DA) techniques to bridge the in *vitro* to in vivo gap [25,26,30].

A shared underlying principle for MTL and DA techniques is that *transfer learning,* whether it is across tasks or domains, needs generalization of information through shared representations. Inability to do this effectively leads to negative transfer (NT) where predictive performance for target tasks or domains is instead hampered relative to single task learning (STL) [37]. For MTL, this can happen when unrelated tasks are learnt together (potentially addressed by quantifying *task relatedness* as in GO-MTL [22]) or when poor predictors adversely impact the shared representation (potentially addressed by weighting transfer flows based on task loss as in AMTL [23] and its extension Deep-AMTFL [24]). For DA, NT can occur when there is weak or no similarity between domains [21] and the method PRECISE [26] seeks to address this for drug response prediction via a robust manifold alignment process. A refinement of this idea, TRANSACT [25], uses Kernel-PCA based sub-space alignment to further capture non-linear relationships between samples from *in vitro* and *in vivo* domains. Existing methods however rely on the covariate-shift assumption, where marginal distributions for features (*P_s_*(*X*) and *P_t_*(*X*), for tasks/domains *s* and *t*) are allowed to vary while the conditional distribution for drug response is assumed to be the same (*P_s_*(*Y*|*X*) = *P_t_*(*Y*|*X*)) [21,39]. This strict assumption can often lead to NT [1,39] when e.g. drugs that are effective *in vitro* do not successfully translate to the clinical setting [36].

We present a unified transfer learning approach (TUGDA) for MTL and DA that leverages task/domain uncertainty (rather that loss) and a relaxed covariate-shift assumption to improve robustness of drug response prediction. Specifically, TUGDA captures both *aleatoric* [19] and *epistemic* [20] uncertainties, and uses them to weight the task/domain to feature transfer. In addition, TUGDA relaxes the covariate-shift assumption across domains (*P_s_*(*Y* |*X*) ≈ *P_t_*(*Y*|*X*)) for tasks with low confidence predictions using shared domain features. Our evaluations against state-of-the-art methods show that the use of uncertainties in guiding task-to-feature transfer reduces cases of negative transfer 94% overall and by 63% in harder cases that have limited *in vitro* data. For *in vivo* settings, TUGDA outperformed previous methods in transferring drug responses to patient-derived xenograft (PDX) models, while providing comparable but complementary performance with patient data. Overall, TUGDA represents a novel unified framework to leverage information from *in vitro* and *in vivo* settings, and robustly predict cancer drug responses from molecular profiles.

## 2 Methods

### 2.1 Definitions and Preliminaries

Similar to prior work [24], we assume a dataset 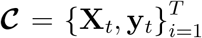 consisting of ***X**_t_* ∈ ℝ^*N_t_*×*d*^ gene expression profiles (*d* genes) and ***y**_t_* ∈ ℝ^*N_t_*×1^ drug response values for *T* different drugs and *N_t_* different cell types (cell lines, xenografts or patients). In a MTL setting, we jointly learn predictive models for all *T* tasks under the following general framework:

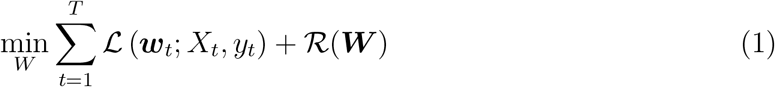

where 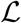 is the loss function (e.g. mean squared error, in our case) applied to each task *t*, with ***w_t_*** representing task-specific parameters as columns of ***W*** ∈ ℝ^*d*×*T*^. The regularization term R is introduced to enforce priors over the task parameters and to improve generalization. This approach constrains joint learning in a naive manner (through the regularization term) and an approach to improve this is to assume that there exist shared latent bases across tasks [3,22]. We can represent this assumption and improve eq. (1) as follows:

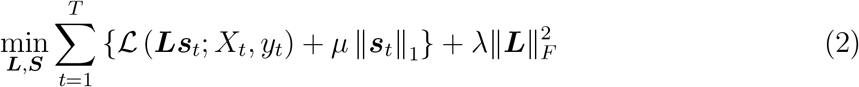

where W from eq. (1) is decomposed as ***W*** = ***LS***, with ***L*** ∈ ℝ^*d*×*k*^ representing the set of *k* latent bases, and ***S*** ∈ ℝ^*k*×*T*^ is the matrix containing vectors ***s_t_*** to combine those bases. The 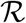 term from eq. (1) is then replaced to constrain ***L*** to be ℓ_2_ regularized while ***s**_t_* needs to be *ℓ*_1_ sparse, with the hyper-parameters *μ* and *σ* controlling the extent of regularization. This framework can be extended to take advantage of neural networks and use multiple layers of shared features followed by a task-specific layer. Here we assume that ***L*** and ***S*** are parameters for the first and the second (task-specific) hidden layers, respectively.

The approach in eq. (2) tries to reduce the risk of negative transfer by forcing unrelated tasks to use disjoint latent spaces. Nevertheless shared bases are trained without consideration of the quality of task-predictors, allowing for noisy and unreliable predictors to be the source of NT [24]. Assuming that task loss is a proxy for task reliability, the transfer from task-to-features can be guided [23,24] by extending eq. (2) as follows:

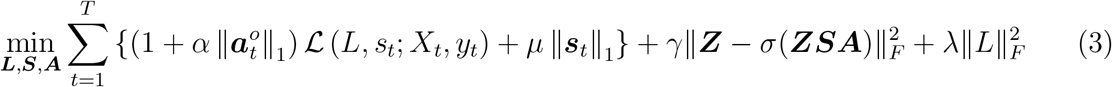

where ***Z*** = *σ*(***X**_t_**L***) is the output of the first neural layer ***L*** followed by a non-linear activation function (ReLU [27] in our case), ***Z*** is interpreted as the shared features space and it is used by ***S*** (task-specific parameters) to predict the drug responses. ***A*** is a matrix which controls the amount of transfer from task *t* to *k* features by the row vector 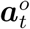 (***A***’s row vector). An auto-encoder regularization is then imposed aiming to reconstruct the latent features ***Z*** with the model output ***ZSA***. Note that the hyperparameter *α* is multiplied by the training loss 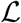 to control the sparsity of 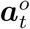, thus breaking the symmetry of transfer to features by forcing transfer from high loss tasks to be more sparse [23,24]. Despite this sophisticated formulation, the assumption that task loss is a proxy for reliability may be misleading, especially in cases of overfitting from limited *in vitro* training data [14].

### 2.2 Leveraging task uncertainty for multi-task learning

Following Kendall et al [19], we aimed to estimate two types of task uncertainties and explored their use as alternative weights for task-to-feature transfer. The first type is *aleatoric* uncertainty which captures uncertainty due to inherent noise in the experimental data that is being modelled. Specifically, as shown by [20] *homoscedastic aleatoric* uncertainty in MTL settings captures the relative confidence between tasks. As this uncertainty does not vary with input data, we can interpret it as task uncertainty reflecting the amount of noise inherent in drug response measurements. Let **f**^w_t_^ (**x**) be the output function for input **x** and task-weight **w**_t_, we have the following relationship for *aleatoric* uncertainty per task (*σ_t_*) in a regression setting:

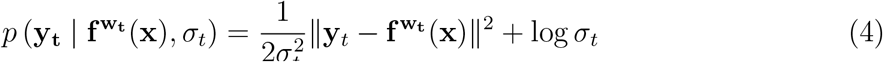

where *σ* is learnable along with model parameters. Intuitively from eq. (4), *σ_t_* can been interpreted as loss attenuation when the model predictions are far away from the ground truth. As prior work has shown that MTL is strongly impacted by relative weighting for task-to-feature transfer [19], the use of *aleatoric* uncertainty in TUGDA could reduce NT by automatically learning optimal weights.

A second type of task uncertainty that is accounted for in TUGDA is *epistemic,* representing the uncertainty in model parameters [19]. To do so, TUGDA uses Bayesian neural networks (BNNs, where weights 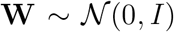) to quantify model prediction uncertainties [13]. We used dropout variational inference for approximate inference in our model [10] using dropout during training [31] and testing, thus enabling sampling from an approximate posterior distribution for weights 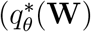, in a tractable family) that minimizes Kullback-Leibler divergence to the true model posterior [10]. We therefore extend eq. (4) as:

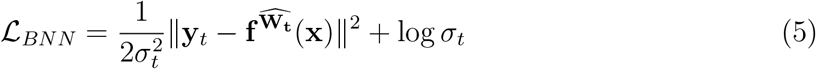

where 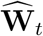 is sampled from the approximate distribution 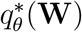. In this setting, predictions are obtained by forwarding each sample *x* though the model for *P* passes, with weights sampled according to dropout inference.

In the TUGDA framework, with BNN feature extractor *F*, BNN latent space *Z*, a multi-task layer *S* and a decoder layer *A* to regularize transfer from task-to-features, the epistemic uncertainty for a task *t* given a sample *x* is computed in *P* passes as:

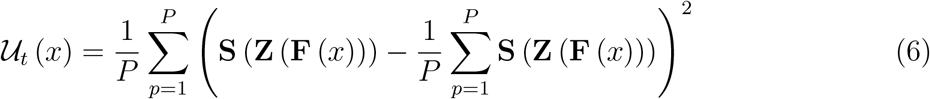

The use of task-uncertainties to guide knowledge transfer from tasks-to-features is then accomplished by:

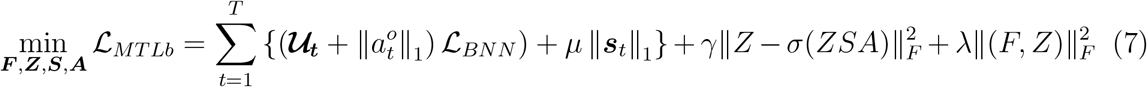

where 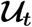 is employed to weight 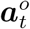, thus forcing tasks with high-uncertainty to transfer less to the shared feature space. A model representation for MTL with TUGDA is depicted in fig. 1 (blue layers) showing how the influence of task-loss is attenuated by both *aleatoric* 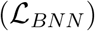 and *epistemic* 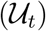 uncertainties, and how constraints for 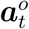 are learnt in an end-to-end fashion.

**Fig. 1.**
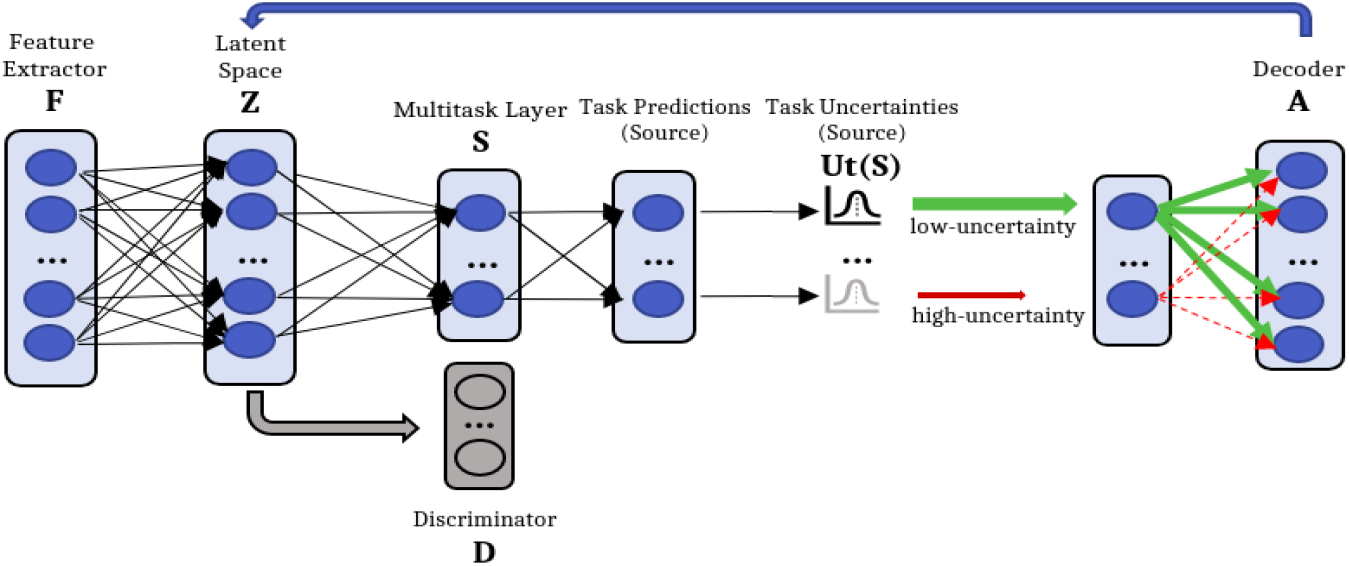
TUGDA framework for multi-task learning and domain adaptation in cancer drug response prediction: The feature extractor F receives input data and maps them to a latent space Z. Then, the multi-task layer S uses latent features to make predictions, as well as compute task-uncertainties 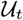 for regularizing the amount of transfer from tasks/domains in A to the latent features in Z. Using adversarial learning, the discriminator D receives the extracted features from Z and regularizes F to learn domain-invariant features. F, Z, S, A and D consist of a single fully connected layer.

### 2.3 Domain Adaptation with task uncertainty and relaxed covariate-shift assumption

To enable domain adaptation from *in vitro* to *in vivo* settings while avoiding NT for tasks where similarity between domains is limited, we extend eq. (7) by adding a Discriminator module *D* (fig. 1, D in gray) that is responsible for classifying a sample *x* into different domains (cell line, xenograft or patient tumor). The idea here is to use adversarial learning to match source (*in vitro*) and target (*in vivo*) marginal distributions [11]. In this manner, we can describe the training process as a two-player game, where the module *Z*(*F*(*x*)) learns features that forces *D*(*Z*(*F*(*x*))) towards confusion, while D needs to accurately classify domains (fig. 1, blue and gray). In the end, *Z*(*F*(*x*)) is expected to learn features that are domain-invariant and so we can describe the learning process as:

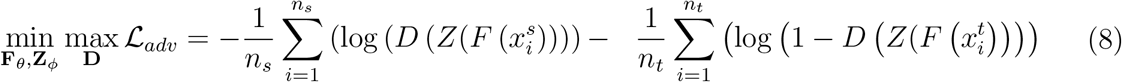

with *n_s_* and *n_t_* being the number of training samples from source (*in vitro*) and target (*in vivo*), respectively. To enable this adversarial training, we employ the Gradient Reversal Layer (GRL) approach [11] that works by flipping the sign of gradients that flow through *D* to the network during back-propagation. With this change, we have a model that jointly learns a shared space (aligns the marginals) and uses these features to predict cancer drug response in a MTL setting. As we regularize transfer from task-to-features using task-uncertainties, we constrain our model to allow predictions with high uncertainty using shared features to transfer less. In contrast to previous methods [25,26], we impose a relaxed covariate-shift for transferring information from high-uncertainty predictors under the assumption that they are less likely to retain predictions across domains. With this TUGDA is trained in an end-to-end fashion as follows:

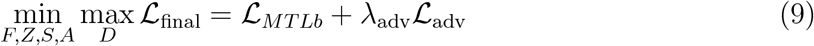

with λ_adv_ as a hyper-parameter which controls the influence of adversarial training.

## 3 Experiments and Results

### 3.1 TUGDA reduces negative transfer in multi-task learning of in *vitro* drug responses

#### Dataset and Baselines

To evaluate the MTL performance of TUGDA (fig. 1, blue), we used the *Genomics of Drug Sensitivity in Cancer* (GDSC) database [16] to obtain cell line drug response and transcriptomic data. Following the steps in Mourragui et al [25] to pre-process data, we obtained a matrix of normalized gene expression values for 806 celllines and 1,780 genes [15], in addition to response values for 200 drugs. As prior work has shown that regularized linear models often yield state-of-the-art results [8,17], we employed ridge linear regression as a single-task learning baseline. We then compared TUGDA with the state-of-the-art neural network based multi-task learners GO-MTL [22]), AMTL [23] and Deep-AMTFL [24]), that are designed to avoid NT behavior. By combining the feature extractor *F* with GO-MTL and AMTL we obtained two additional baselines that we refer to as Deep-GO-MTL and Deep-AMTL, respectively. Thus, all deep neural network models share the same number of layers until the prediction step (F, Z and S; fig. 1), and the differences are only in terms of the regularization used. We performed 3-fold cross-validation to tune hyper-parameters for all methods and used the *Tree-structured Parzen Estimator* algorithm [5] to search for the best set of hyper-parameters (list and range in supplement S1).

#### Results

Models were trained to predict IC50 values (concentration which kills 50% of cells; log-transformed) and compared in terms of mean squared error (MSE) distribution across all 200 drugs (3-fold cross-validation). As can be seen in fig. 2a, TUGDA improves over prior methods with the lowest mean MSE of 1.80. Higher performances were also observed in our ablation analysis, which consists of the following setup: TUGDA(A) is solely based on *aleatoric* uncertainty; TUGDA(E) uses *epistemic* uncertainty; and TUGDA(E+A) uses both uncertainty types but excludes the use of *U_t_* to weight 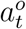. This analysis suggests that epistemic uncertainty is key to TUGDA’s performance in this dataset.

**Fig. 2.**
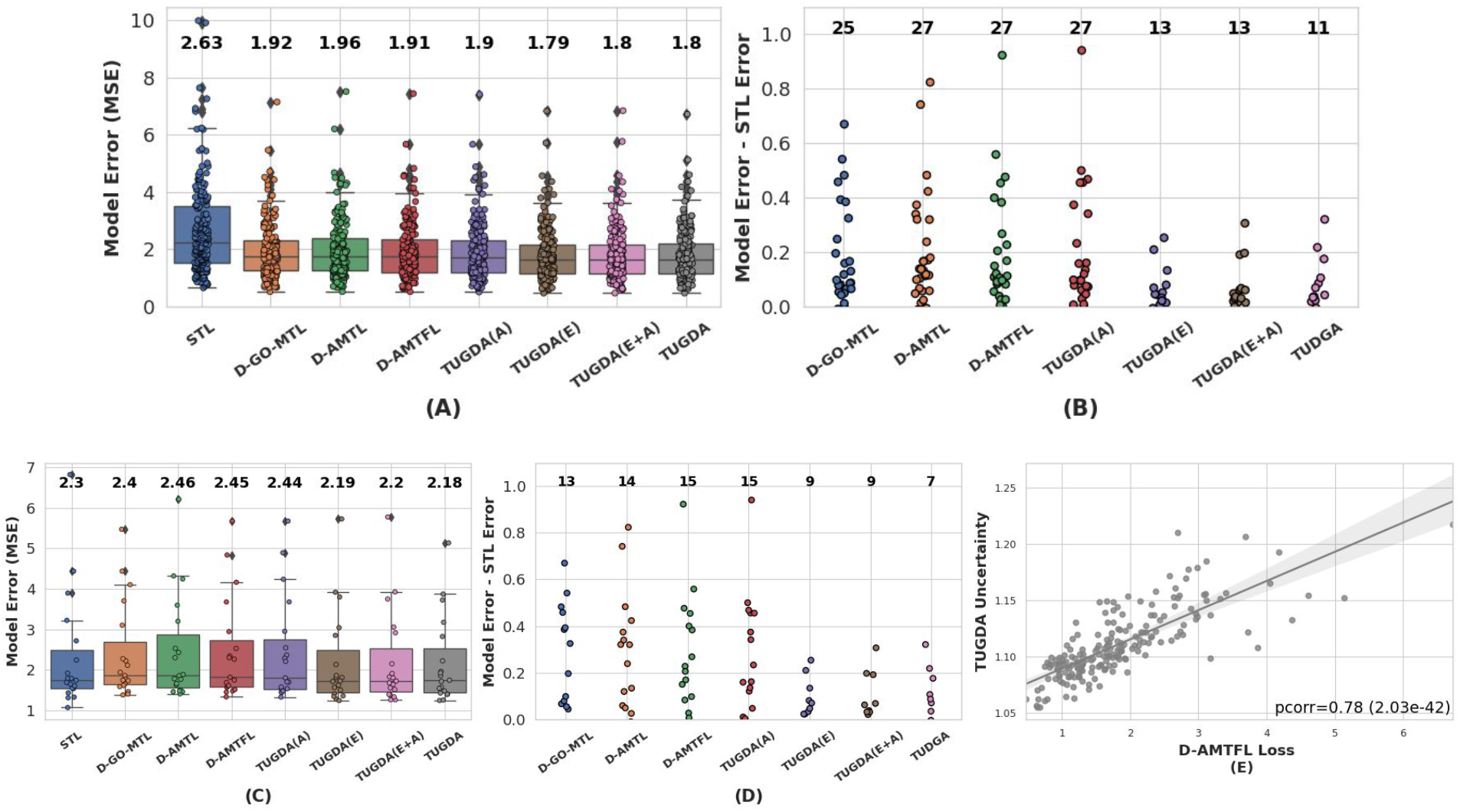
MTL performance evaluation using *in vitro* datasets. (a) Boxplots showing MSE across tasks for different models including STL (baseline), state-of-the-art methods (Deep-GO-MTL, Deep-AMTL, Deep-AMTFL), and TUGDA and its ablated variantss (mean MSE values are shown on the top), (b) Strip plots comparing the degree of negative transfer and the number of such tasks (shown on top). (c) and (d) Boxplots and strip plots comparing MSE and NT for tasks with smaller sample sizes (n=19). (e) Scatter plot comparing Deep-AMTFL’s task-loss and TUGDA’s task-uncertainty values.

To quantify NT behavior, STL-based MSEs were subtracted from corresponding MTL-based MSEs s.t. positive values indicate NT. As shown in fig. 2b, TUGDA presented the fewest number of NT cases (11 out of 200 tasks, 94% of tasks with no NT), reducing the number of tasks with NT by 56% for the next best method (Deep-GO-MTL). We next focused our analysis of performance on the more challenging tasks with smaller sample sizes (n=19; median = 49, max = 382) compared to the rest (sample size min = 716, median = 745). In this setting, TUGDA outperformed existing methods in terms of mean MSE (fig. 2c), and the ablation analysis highlights the utility of the full model. As can be seen from fig. 2d, NT cases are clearly enriched in this set of 19 tasks and TUGDA reduces the number from 13 (for the next best method, Deep-GO-MTL) to 7 (63% of tasks without NT; reduction of NT tasks by 46% relative to Deep-GO-MTL). Overall, Deep-AMTFL’s task-loss and TUGDA’s task-uncertainties were observed to be correlated to some extent (Pearson r = 0.78), though there were differences for some tasks indicating the spots where TUGDA’s can improve overs task-loss based MTL fig. 2e).

### 3.2 TUGDA provides a robust approach for domain adaptation from *in vitro* to *in vivo* response prediction

#### Datasets and Baselines

We evaluated the unified TUGDA framework (fig. 1, blue and gray modules) against existing unsupervised domain adaptation (UDA) methods for transferring cancer drug responses from cell-lines (*in vitro*) to two different *in vivo* settings, patient-derived xenografts (PDX) and patient tumors. PDX data was obtained from the Novartis Institutes for Biomedical Research [12], containing gene expression profiles (n = 399) and drug responses values. Patient tumor expression profiles were obtained from TCGA [28] as well as curated response data from Ding et al [9]. All cell line, PDX and tumor data were processed using the same pipelines, and the same pre-processing steps and experimental setup as proposed in Mourragui et al [25]. An elastic net regression trained solely on cell line data as well the UDA approaches PRECISE [26] and TRANSACT[25] served as baselines for both PDX and patient tumor predictions. For PDX data, TUGDA was trained following the approach in Ganin et al [11] where the set of hyperparameters was determined by minimizing MSE loss on test-fold data (cell line). For patient tumors, hyper-parameters were selected to maximize spearman correlation of cell-line and PDX response values. In both cases, the *Tree-structured Parzen Estimator* algorithm [5] was used for searching hyper-parameters (list and range in supplement S2).

#### Results with PDX data

We evaluated the transfer of drug responses from GDSC celllines to PDX data based on 12 shared drugs (extended from [25]) and computed spearman correlations for predicted and measured response values in the PDX setting (PDX best average response). Out of 12 drugs, TUGDA provided the highest spearman correlation for 6 drugs (fig. 3a, Buparlisib, Cetuximab, LGK974, Paclitaxel, Tamoxifen and Trametinib), while TRANSACT, PRECISE and Elastic Net were the best methods for 5, 0 and 1 drugs, respectively. Investigating the learnt feature space, we observed that cell-lines and PDX samples from the same tissue tend to cluster together, showing that the model infers a biologically appropriate *in vitro* to *in vivo* transformation (fig. 3b).

**Fig. 3.**
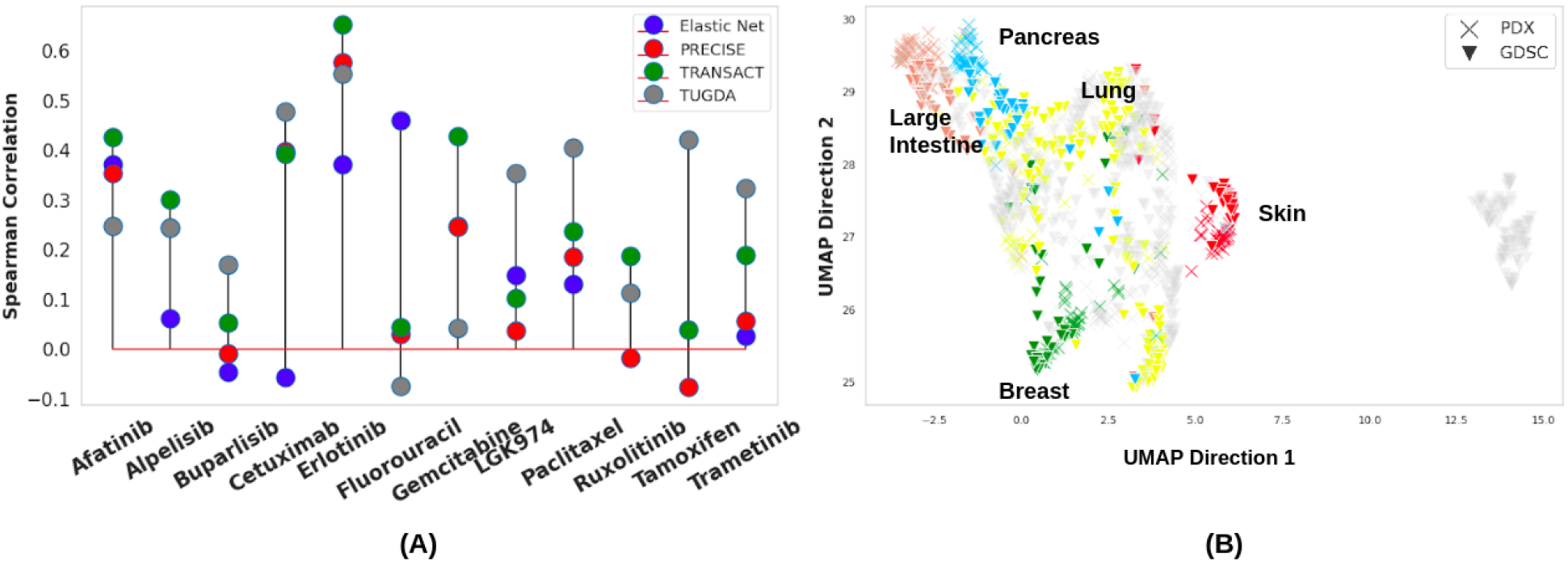
DA performance for predicting drug response in PDX models. (a) Comparison of Spearman correlation between cell-line and PDX response values for 12 drugs across different models. (b) UMAP plot of the feature space from TUGDA showing clusters of cell-line and PDX samples that originate from the same tissue. Gray points have undefined tissue types.

#### Results with patient data

For patient tumor data, we evaluated performance for transferring drug response predictions from cell-lines to patients based on 22 drugs shared in GDSC and TCGA (extended from [25]). As analyzed previously [25,9], TCGA drug responses were categorized into two groups, *Responders* (“Complete Response” and “Partial Response”) and *Non-responders* (“Stable Disease” and ‘Progressive Disease”). Despite several additional sources of variation in patient response data (tumor heterogeneity and environment, immune response, patient health status), TUGDA showed significant associations for 7 drugs including Bleomycin, Docetaxel, Doxorubicin, Gemcitabine, Paclitaxel, Premetexed and Tamoxifen (Table 1; one-sided Mann-Whitney test between predicted responses for Responders and Non-responders; p-value < 0.05), improving on PRECISE (6 drugs) and Elastic Net (3 drugs) baselines, and complementing predictions from TRANSACT (7 drugs) with significant association in 5 additional drugs. As was the case for the PDX model, the UMAP projection of the learnt feature space from TUGDA largely clusters cell-line and patient tumor data by tissue type (fig. 4), highlighting that it can successfully learn shared biological properties.

**Fig. 4.**
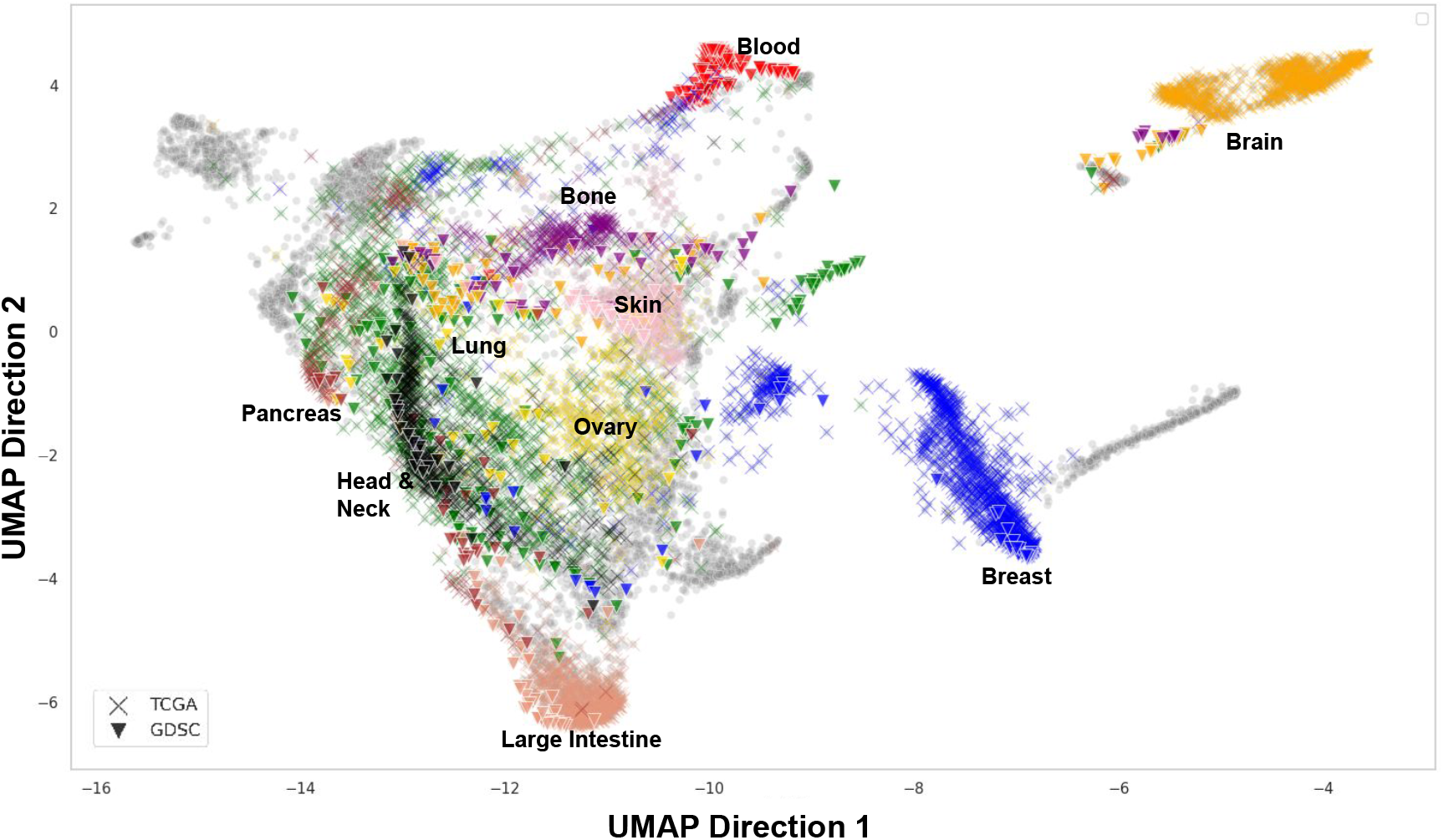
UMAP plot of feature space from TUGDA shows tissue-type specific clusters of cell-line and patient tumor data.

**Table 1.** DA performance for predicting drug response in patient data. Drug names in parenthesis are corresponding matches from GDSC.

| Drug | Samples | Elastic Net | PRECISE | TRANSACT | TUGDA |
| --- | --- | --- | --- | --- | --- |
| Bicatulamide | 17 | 0.201 | 0.116 | 0.244 | 0.285 |
| Bleomycin | 53 | 0.153 | 0.082 | 0.091 | **0.009** |
| Cetuximab | 19 | 0.119 | 0.484 | 0.076 | 0.196 |
| Cisplatin | 308 | **1.10e-6** | **2.14e-5** | **7.15e-7** | 0.616 |
| Carboplatin (Cisplatin) | 166 | **0.042** | **0.024** | **0.003** | 0.0701 |
| Cyclophosphamide | 102 | 0.571 | 0.112 | 0.587 | 0.598 |
| Dacarbazine (AICAR) | 30 | 0.323 | 0.385 | 0.692 | 0.1206 |
| Docetaxel | 102 | 0.617 | 0.762 | 0.111 | **1.12e-4** |
| Doxorubicin | 101 | 0.805 | 0.998 | 0.702 | **1.15e-06** |
| Epirubicin | 25 | 0.451 | 0.678 | 0.19 | 0.546 |
| Etoposide | 84 | 0.201 | **0.007** | **0.025** | 0.81 |
| 5-Fluorouracil | 186 | 0.219 | 0.8 | 0.361 | 0.327 |
| Gemcitabine | 156 | 0.248 | **0.039** | **0.003** | **0.02** |
| Irinotecan | 25 | 0.81 | 0.536 | 0.463 | 0.892 |
| Oxaliplatin | 66 | 0.379 | **0.027** | **0.035** | 0.991 |
| Paclitaxel | 160 | **0.032** | 0.129 | **0.004** | **0.013** |
| Premetexed | 38 | 0.157 | 0.178 | 0.189 | **0.018** |
| Tamoxifen | 23 | 0.999 | 0.487 | 0.895 | **0.038** |
| Temozolomide | 96 | 0.555 | 0.586 | 0.182 | 0.543 |
| Trastuzumab (Afatinib) | 16 | 0.089 | **0.024** | **0.016** | 0.532 |
| Vinblastine | 16 | 0.298 | 0.5 | 0.583 | 0.375 |
| Vinorelbine | 30 | 0.312 | 0.163 | 0.384 | 0.754 |

## 4 Conclusion

TUGDA’s strength lies in the fact that it represents a unified transfer learning approach for multi-task learning and domain adaptation that leverages the concept of task/domain uncertainty. These attributes align it to the fundamental challenges found in building predictive models for precision oncology, including sample size limitations, lack of curated *in vivo* data and violations of the covariate-shift assumption. Our experiments show that TUGDA can provide notable benefits in a multi-task setting to reduce negative transfer, particularly when training data is limited. In addition, it shows promise as a way to robustly transfer information from *in vitro* data to *in vivo* settings, based on confidence in task predictions. This is aided by many neural network features that help avoid overfitting, including the use of Bayesian neural networks, L1 and L2 regularizations for feature and task-specific layers, dropouts, and the incorporation of task-uncertainties for regularizing task-to-feature transfer (instead of attention weights [29]).

TUGDA’s approach to relaxing the covariate-shift assumption is a natural by-product of MTL in a adversarial domain adaptation framework. This is distinct from prior work [2] that is based on learning the probability of label changes across source and target domains and using this to weight transfer. In a recent study, the concept of *label-shift* has also been highlighted as a source of NT [33]. Intrinsic differences in cancer cell lines and patient tumors (e.g. the enrichment of genomic alterations and *in vitro* selection of subpopulations) make this scenario a likely one for domain adaptation in precision oncology. We envisage that TUGDA’s framework can be extended to alleviate NT in the marginal distribution of drug responses as well, advancing the goal of realistic precision oncology models further.

## Supporting information

Hyperparameters tested.

## References

1. Latent-variable models for drug response prediction and genetic testing (2020), https://tspace.library.utoronto.ca/handle/1807/100951

2. Adel, T., Zhao, H., Wong, A.: Unsupervised domain adaptation with a relaxed covariate shift assumption. In: Proceedings of the Thirty-First AAAI Conference on Artificial Intelligence. p. 1691—1697. AAAI’17, AAAI Press (2017)

3. Argyriou, A., Evgeniou, T., Pontil, M.: Convex multi-task feature learning. Machine Learning 73(3), 243—272 (Dec 2008). https://doi.org/10.1007/s10994-007-5040-8, https://doi.org/10.1007/s10994-007-5040-8

4. Azuaje, F.: Computational models for predicting drug responses in cancer research. Briefings in Bioinformatics 18(5), 820—829 (07 2016). https://doi.org/10.1093/bib/bbw065, https://doi.org/10.1093/bib/bbw065

5. Bergstra, J., Bardenet, R., Bengio, Y., Kégl, B.: Algorithms for hyper-parameter optimization. In: Proceedings of the 24th International Conference on Neural Information Processing Systems. p. 2546—2554. NIPS’11, Curran Associates Inc., Red Hook, NY, USA (2011)

6. Brown, N.A., Elenitoba-Johnson, K.S.: Enabling precision oncology through precision diagnostics. Annual Review of Pathology: Mechanisms of Disease 15(1), 97—121 (2020). https://doi.org/10.1146/annurev-pathmechdis-012418-012735, https://doi.org/10.1146/annurev-pathmechdis-012418-012735, pMID: 31977297

7. Chae, Y.K., Pan, A.P., Davis, A.A., Patel, S.P., Carneiro, B.A., Kurzrock, R., Giles, F.J.: Path toward precision oncology: Review of targeted therapy studies and tools to aid in defining “actionability” of a molecular lesion and patient management support. Molecular Cancer Therapeutics 16(12), 2645—2655 (2017). https://doi.org/10.1158/1535-7163.MCT-17-0597, https://mct.aacrjournals.org/content/16/12/2645

8. Costello, J., Community, N.C.I.D.R.E.A.M.: A community effort to assess and improve drug sensitivity prediction algorithms. Nature Biotechnology 32(12), 1202—1212 (Dec 2014). https://doi.org/10.1038/nbt.2877, https://doi.org/10.1038/nbt.2877

9. Ding, Z., Zu, S., Gu, J.: Evaluating the molecule-based prediction of clinical drug responses in cancer. Bioinformatics 32(19), 2891—2895 (06 2016). https://doi.org/10.1093/bioinformatics/btw344, https://doi.org/10.1093/bioinformatics/btw344

10. Gal, Y., Ghahramani, Z.: Dropout as a bayesian approximation: Representing model uncertainty in deep learning. In: Proceedings of the 33rd International Conference on International Conference on Machine Learning - Volume 48. p. 1050—1059. ICML’16, JMLR.org (2016)

11. Ganin, Y., Lempitsky, V.: Unsupervised domain adaptation by backpropagation. In: Proceedings of the 32nd International Conference on International Conference on Machine Learning - Volume 37. p. 1180—1189. ICML’15, JMLR.org (2015)

12. Gao, H.e.a.: High-throughput screening using patient-derived tumor xenografts to predict clinical trial drug response. Nature Medicine 21(11), 1318–1325 (Nov 2015). https://doi.org/10.1038/nm.3954, https://doi.org/10.1038/nm.3954

13. Goan, E., Fookes, C.: Bayesian neural networks: An introduction and survey p. 45—87 (2020)

14. Hawkins, D.M.: The problem of overfitting. Journal of Chemical Information and Computer Sciences 44(1), 1—12 (2004). https://doi.org/10.1021/ci0342472, https://doi.org/10.1021/ci0342472, pMID: 14741005

15. Hoogstraat, M., de Pagter, M.S., Cirkel, G.A., van Roosmalen, M.J., Harkins, T.T., Duran, K., Kreeftmeijer, J., Renkens, I., Witteveen, P.O., Lee, C.C., Nijman, I.J., Guy, T., van ’t Slot, R., Jonges, T.N., Lolkema, M.P., Koudijs, M.J., Zweemer, R.P., Voest, E.E., Cuppen, E., Kloosterman, W.P.: Genomic and transcriptomic plasticity in treatment-naïve ovarian cancer. Genome Research 24(2), 200—211 (2014). https://doi.org/10.1101/gr.161026.113

16. Iorio, F.D., Knijnenburg, T.A., Vis, D.J., Bignell, G.R., Menden, M.P., Schubert, M.B., Aben, N., Gonçalves, E., Barthorpe, S., Lightfoot, H., Cokelaer, T., Greninger, P., van Dyk, E., Chang, H.C., de Silva, H., Heyn, H., Deng, X., Egan, R.K., Liu, Q., Mironenko, T., Mitropoulos, X., Richardson, L.B., Wang, J., Zhang, T., Moran, S., Sayols, S., Soleimani, M., Tamborero, D., López-Bigas, N., Ross-MacDonald, P., Esteller, M., Gray, N.S., Haber, D.A., Stratton, M.R., Benes, C.H., Wessels, L.F.A., Saez-Rodriguez, J., McDermott, U., Garnett, M.J.: A landscape of pharmacogenomic interactions in cancer. In: Cell (2016)

17. Jang, I.S., Neto, E.C., Guinney, J., Friend, S.H., Margolin, A.A.: Systematic assessment of analytical methods for drug sensitivity prediction from cancer cell line data. Pacific Symposium on Biocomputing. Pacific Symposium on Biocomputing pp. 63—74 (2014), https://pubmed.ncbi.nlm.nih.gov/24297534, 24297534[pmid]

18. Jiang, P., Sellers, W.R., Liu, X.S.: Big data approaches for modeling response and resistance to cancer drugs. Annual Review of Biomedical Data Science 1(1), 1—27 (2018). https://doi.org/10.1146/annurev-biodatasci-080917-013350, https://doi.org/10.1146/annurev-biodatasci-080917-013350

19. Kendall, A., Gal, Y.: What uncertainties do we need in bayesian deep learning for computer vision? (2017)

20. Kendall, A., Gal, Y., Cipolla, R.: Multi-task learning using uncertainty to weigh losses for scene geometry and semantics (2018)

21. Kouw, W.M., Loog, M.: A review of domain adaptation without target labels (2019)

22. Kumar, A., au2, H.D.I.: Learning task grouping and overlap in multi-task learning (2012)

23. Lee, G., Yang, E., Hwang, S.J.: Asymmetric multi-task learning based on task relatedness and loss. In: Proceedings of the 33rd International Conference on International Conference on Machine Learning - Volume 48. p. 230—238. ICML’16, JMLR.org (2016)

24. Lee, H.B., Yang, E., Hwang, S.J.: Deep asymmetric multi-task feature learning (2018)

25. Mourragui, S., Loog, M., Vis, D.J., Moore, K., Manjon, A.G., van de Wiel, M.A., Reinders, M.J., Wessels, L.F.: Precise+ predicts drug response in patients by non-linear subspace-based transfer from cell lines and pdx models. bioRxiv (2020). https://doi.org/10.1101/2020.06.29.177139, https://www.biorxiv.org/content/early/2020/07/28/2020.06.29.177139

26. Mourragui, S., Loog, M., van de Wiel, M.A., Reinders, M.J.T., Wessels, L.F.A.: PRECISE: a domain adaptation approach to transfer predictors of drug response from pre-clinical models to tumors. Bioinformatics 35(14), i510—i519 (07 2019). https://doi.org/10.1093/bioinformatics/btz372, https://doi.org/10.1093/bioinformatics/btz372

27. Nair, V., Hinton, G.E.: Rectified linear units improve restricted boltzmann machines. In: Proceedings of the 27th International Conference on International Conference on Machine Learning. p. 807—814. ICML’10, Omnipress, Madison, WI, USA (2010)

28. Network, C.G.A.R., Weinstein, J.N., Collisson, E.A., Mills, G.B., Shaw, K.R.M., Ozenberger, B.A., Ellrott, K., Shmulevich, I., Sander, C., Stuart, J.M.: The cancer genome atlas pan-cancer analysis project. Nature genetics 45(10), 1113—1120 (Oct 2013). https://doi.org/10.1038/ng.2764, https://pubmed.ncbi.nlm.nih.gov/24071849, 24071849[pmid]

29. Nguyen, T.A., Jeong, H., Yang, E., Hwang, S.J.: Clinical risk prediction with temporal probabilistic asymmetric multi-task learning (2020)

30. Sharifi-Noghabi, H., Peng, S., Zolotareva, O., Collins, C.C., Ester, M.: AITL Adversarial Inductive Transfer Learning with input and output space adaptation for pharmacogenomics. Bioinformatics

31. Srivastava, N., Hinton, G., Krizhevsky, A., Sutskever, I., Salakhutdinov, R.: Dropout: A simple way to prevent neural networks from overfitting. Journal of Machine Learning Research 15(56), 1929—1958 (2014), http://jmlr.org/papers/v15/srivastava14a.html

32. Suphavilai, C., Bertrand, D., Nagarajan, N.: Predicting cancer drug response using a recommender system. Bioinformatics (2018)

33. Tan, S., Peng, X., Saenko, K.: Class-imbalanced domain adaptation: An empirical odyssey (2020)

34. van Staveren, W., Solís, D.W., Hébrant, A., Detours, V., Dumont, J., Maenhaut, C.: Human cancer cell lines: Experimental models for cancer cells in situ? for cancer stem cells? Biochimica et Biophysica Acta (BBA) - Reviews on Cancer 1795(2), 92—103 (2009). https://doi.org/10.1016/j.bbcan.2008.12.004, http://www.sciencedirect.com/science/article/pii/S0304419X08000796

35. Wang, L., Li, X., Zhang, L., Gao, Q.: Improved anticancer drug response prediction in cell lines using matrix factorization with similarity regularization. BMC Cancer 17(1), 513 (Aug 2017). https://doi.org/10.1186/s12885-017-3500-5, https://doi.org/10.1186/s12885-017-3500-5

36. Wilding, J.L., Bodmer, W.F.: Cancer cell lines for drug discovery and development. Cancer Research 74(9), 2377—2384 (2014). https://doi.org/10.1158/0008-5472.CAN-13-2971, https://cancerres.aacrjournals.org/content/74/9/2377

37. Zhang, W., Deng, L., Wu, D.: Overcoming negative transfer: A survey (2020)

38. Zhang, Y., Yang, Q.: A survey on multi-task learning (2018)

39. Zhao, H., des Combes, R.T., Zhang, K., Gordon, G.J.: On learning invariant representation for domain adaptation (2019)

