## Supplementary material for "TUGDA: Task uncertainty guided domain adaptation for robust generalization of cancer drug response prediction from *in vitro* to *in vivo* settings": Hyperparameters tested.

In MTL settings we performed 3-fold cross-validation to tune hyper-parameters for all methods and used the *Tree-structured Parzen Estimator* algorithm [5] to search for the best set of hyper-parameters.

### S1 - Hyperparameters tested on MTL experiments

We searched for the best set parameters for each model in the following space:

F layer Units = 64, 128, 256, 512, 1024
Z layer Units = 200, 300, 400, 500, 600, 700
Learning Rates = 1e-2, 1e-3, 1e-4, 1e-5, 5e-2, 5e-3, 5e-4, 5e-5
Mu = 1, 0.1, 0.01, 0.001, 0.0001
Lambda = 1, 0.1, 0.01, 0.001, 0.0001
Gamma = 1, 0.1, 0.01, 0.001, 0.0001
Alpha = 1, 0.1, 0.01, 0.001, 0.0001
Dropout = 0.1, 0.2, 0.3, 0.4, 0.5

For all epistemic and TUGDA models we sampled 50 times to compute the task-uncertainties. All epochs were fixed (100 epochs). Batch size is fixed at 300 to keep all tasks containing at least one sample each from each task. The Layer A in charge for reconstructing the latent space Z contains the same number of units as Z.

#### Best settings found for each model

1) Deep-GO-MTL
F layer Units = 64, Z layer Units = 200, Lr= 0.01, Mu= 0.0001, Lambda= 0.01

2) Deep-AMTL
F layer Units = 64 , Z and A layer Units = 700, Lr= 0.01, Mu= 0.001, Lambda= 1, Alpha= 0.001, Gamma= 0.0001

3) Deep-AMTFL
F layer Units = 128, Z and A layer Units = 700, Lr= 0.01, Mu= 0.01, Lambda= 1, Alpha= 0.001, Gamma= 0.0001

4) TUGDA (A)

F layer Units = 128, Z and A layer Units = 700, Lr= 0.01, Mu= 0.01, Lambda= 1, Alpha= 0.001, Gamma= 0.0001

5) TUGDA (E)
F layer Units = 1024, Z and A layer Units = 700, Lr= 0.005, Mu= 0.001, Lambda= 1, Alpha= 0.1, Gamma= 0.001, Dropout = 0.2

6) TUGDA (E+A)

F layer Units = 1024, Z and A layer Units = 700, Lr= 0.005, Mu= 0.0001, Lambda= 1, Alpha= 0.1, Gamma= 0.001, Dropout = 0.2

7) TUGDA

F layer Units = 1024, Z and A layer Units = 700, Lr= 0.001, Mu= 0.01, Lambda= 0.001, Gamma= 0.0001, Dropout = 0.1

### S2 - Hyperparameters tested on DA experiments

We extended the search to include the number of units in the layers of the domain discriminator D, lambda disc (the hyperparameter controlling the adversarial learning influence) and the discriminator batch size. As we are handling additional datasets, we extended the search space to increase the model capacity.

**PDX search space**

F layer Units = 512, 1024, 1500
Z layer Units = 800, 900, 1000
Learning Rates = 1e-2, 1e-3, 1e-4, 1e-5, 5e-2, 5e-3, 5e-4, 5e-5
Mu = 1, 0.1, 0.01, 0.001, 0.0001
Lambda = 1, 0.1, 0.01, 0.001, 0.0001
Gamma = 1, 0.1, 0.01, 0.001, 0.0001
Alpha = 1, 0.1, 0.01, 0.001, 0.0001
Dropout = 0.1, 0.2, 0.3, 0.4, 0.5
Lamda disc = 0.1, 0.2, 0.3, 0.4, 0.5, 0.6, 0.7, 0.8, 0.9, 1.0
Batch size disc = 32, 64, 128, 256, 300
Epochs = 20, 30, 40, 50
Discriminator units = 400, 500, 600, 700

**Best settings on PDX**

F layer Units = 1500, Z and A layer Units = 900, Lr= 0.001, Mu= 0.001, Lambda= 0.01, Gamma= 0.0001, Dropout = 0.3, Discriminator units=400, Epochs = 50, Batch size= 32, Lamba disc = 0.8

**TCGA search space**

F layer Units = 1024, 1250, 1500
Z layer Units = 800, 900, 1000
Learning Rates = 1e-2, 1e-3, 1e-4, 1e-5, 5e-2, 5e-3, 5e-4, 5e-5
Mu = 1, 0.1, 0.01, 0.001, 0.0001
Lambda = 1, 0.1, 0.01, 0.001, 0.0001
Gamma = 1, 0.1, 0.01, 0.001, 0.0001
Alpha = 1, 0.1, 0.01, 0.001, 0.0001
Dropout = 0.1, 0.2, 0.3, 0.4, 0.5
Lamda disc = 0.1, 0.2, 0.3, 0.4, 0.5, 0.6, 0.7, 0.8, 0.9, 1.0
Batch size disc = 512, 1024
Epochs = 20, 30, 40, 50
Discriminator units = 500, 600, 700

**Best settings on TCGA**

F layer Units = 1024, Z and A layer Units = 800, Lr= 0.005, Mu= 0.01, Lambda= 0.01, Gamma= 1, Dropout = 0.5, Discriminator units=500, Epochs = 30, Batch size= 1024, Lamba disc = 0.7

Elastic net, PRECISE and TRANSACT results were reproduced following "PRECISE+ predicts drug response in patients by non-linear subspace-based transfer from cell lines and PDX models" https://www.biorxiv.org/content/10.1101/2020.06.29.177139v2.full
